# Mind the translational gap: human microglia differ from mouse microglia in their regulation of K_v_ and K_ir_2.1 channels

**DOI:** 10.64898/2026.03.10.710589

**Authors:** Simone Schilling, Jessica Felk, Majed Kikhia, Alice Podestà, Johanna Hintze, Pawel Fidzinski, Martin Holtkamp, Julia Onken, Thomas Sauvigny, Thilo Kalbhenn, Matthias Simon, Helmut Kettenmann, Matthias Endres, Ria Göttert, Karen Gertz

**Affiliations:** Charité – Universitätsmedizin Berlin, Corporate Member of Freie Universität Berlin and Humboldt-Universität zu Berlin, Department of Neurology and Experimental Neurology, Charitéplatz 1, 10117 Berlin, Germany; Charité – Universitätsmedizin Berlin, Corporate Member of Freie Universität Berlin and Humboldt-Universität zu Berlin, Center for Stroke Research Berlin (CSB), Charitéplatz 1, 10117 Berlin, Germany; German Centre for Cardiovascular Research (DZHK), Partner Site Berlin, Berlin, Germany; Berlin Institute of Health at Charité - Universitätsmedizin Berlin, Charitéplatz 1, 10117 Berlin, Germany; Charité – Universitätsmedizin Berlin, Corporate Member of Freie Universität Berlin and Humboldt-Universität zu Berlin, Einstein Center for Neurosciences Berlin, Charitéplatz 1, 10117 Berlin, Germany; Epilepsy-Center Berlin-Brandenburg, Ev. Krankhaus Königin Elisabeth Herzberge, Herzbergstr. 79, 10365 Berlin, Germany; Charité – Universitätsmedizin Berlin, Corporate Member of Freie Universität Berlin and Humboldt-Universität zu Berlin, Department of Clinical and Experimental Epileptology, Charitéplatz 1, 10117 Berlin, Germany; Charité – Universitätsmedizin Berlin, Corporate Member of Freie Universität Berlin and Humboldt-Universität zu Berlin, Institute of Neurophysiology, Charitéplatz 1, 10117 Berlin, Germany; Charité – Universitätsmedizin Berlin, Corporate Member of Freie Universität Berlin and Humboldt-Universität zu Berlin, Department of Neurosurgery, Charitéplatz 1, 10117 Berlin, Germany; Department of Neurosurgery, University Medical Center Hamburg-Eppendorf, Hamburg, Germany; Department of Neurosurgery, Evangelisches Klinikum Bethel, Universitätsklinikum Ostwestfalen-Lippe, Bielefeld, Germany; Max-Delbrück-Center (MDC) for Molecular Medicine in the Helmholtz Association, Berlin-Buch, Germany; Shenzhen University of Advanced Technology, Shenzhen, China

## Abstract

K^+^ channels are important for controlling membrane potential and regulating functional properties of microglia. Whereas the inward-rectifying K^+^ (K_ir_) channel 2.1 modulates proliferation, voltage-gated K^+^ channels (K_v_) are linked to inflammatory response in mouse microglia (mMG). These channels serve as possible drug targets but little is known regarding their activity in human microglia.

We used patch-clamp recording to study membrane currents of primary human microglia (hMG) and human induced pluripotent stem cell-derived microglia-like cells (hiPSC-MGL) and compared them with mMG. Unlike mMG, hMG and hiPSC-MGL exhibited K_ir_2.1 currents only after LPS+IFN-γ stimulation. Interestingly, K_v_ currents were not observed in hMG or hiPSC-MGL under any condition. While mMG had a progressively ameboid morphology after stimulation, hMG showed few morphological changes and hiPSC-MGL increased ramification.

Overall, the activity of K_ir_2.1 and K_v_ channels in hMG and hiPSC-MGL differs fundamentally from mMG. Our findings highlight differences between species and underscore the need for translational approaches.

## Introduction

Microglia are the resident immune cells of the central nervous system (CNS) and play essential roles in maintaining homeostasis, responding to injury, and regulating neuroinflammation [1]. In their surveillant state, microglia exhibit a highly ramified morphology, continuously monitoring the brain microenvironment *in vivo*. Upon activation, they undergo rapid morphological changes, transitioning to an amoeboid shape characterized by retracted processes and enlarged soma - a phenomenon well-characterized in rodent models [2–4]. This morphological shift is accompanied by functional changes, including the release of cytokines and chemokines such as TNF-α, IL-6, IL-1β, and CCL2 [5]. These secretion patterns are well established in rodent microglia and increasingly reported in human microglial models [6].

Furthermore, microglia express a range of ion channels, which are essential for maintaining membrane potential and regulating context-dependent responses [7]. Patch-clamp electrophysiology allows high-temporal resolution of microglial ion channel activity, under resting and activated conditions, offering insights into microglial responses to pathological stimuli [8]. Rodent microglia have been characterized using patch-clamp recordings under various conditions. *In vivo,* mouse microglia express low levels of K^+^ channels, however, after a pathological challenge, inward-rectifying K^+^ channels are upregulated first [4, 8]. Cultured mouse microglia (mMG) constitutively express these inward-rectifying K^+^ channel. Upon stimulation, e.g. by facial axotomy, stab wound or in the context of stroke, mouse microglia express outward-rectifying, voltage-gated K^+^ channels [4, 9, 10].

K_ir_2.1 (*KCNJ2* gene) is the predominant inward-rectifying K^+^ channel in microglia and contributes to the maintenance of the resting membrane potential, thereby regulating activation and responsiveness. It plays a key role in stabilizing microglial homeostasis and modulating their response to inflammation [11]. Microglia also express several outward-rectifying, voltage-gated K^+^ channels of the K_v_ family, with K_v_1.3 being the most extensively studied [12–14]. K_v_1.3 is a voltage-gated K^+^ channel that contributes to membrane repolarization and calcium influx required for functions such as cytokine release, proliferation, and migration [13]. K_v_1.3 is upregulated in activated mouse microglia *in vivo* and *in vitro*, highlighting its relevance in neuroinflammatory and neurodegenerative conditions [12, 14]. It has therefore been suggested as a potential drug target to alter microglia activity [15]. Although the presence of K_v_1.3 has been reported in human microglia in histology and sequencing data, there is a lack of data demonstrating its functional relevance in human microglia [16].

Translation of microglial findings from mouse models to clinical applications has been challenging, in part due to limited access to human cells and tissue. Primary human microglia in culture (hMG) are a precious resource, as surgical tissue is not widely available and often represents pathological conditions. We have previously studied hMG to investigate the properties of human microglia [17]. In addition, human induced pluripotent stem cell-derived microglia-like cells (hiPSC-MGL) represent an increasingly used alternative model that may better reflect human microglial physiology while reducing reliance on animal experiments [17, 18]. However, their application depends on thorough characterization, and electrophysiological data remains scarce.

In this study, we investigated the properties of hMG and hiPSC-MGL in comparison to the well-established mMG, focusing in particular on electrophysiological functions and the activity of potassium channels. Our work underscores the importance of translational approaches.

## Material and Methods

### Primary mouse microglia (mMG)

All experimental procedures were approved by the respective official committees and carried out in strict accordance with the Animal Welfare Act and the ARRIVE (Animals in Research: Reporting In Vivo Experiments) guidelines.

Primary mouse microglia (mMG) were isolated from C57BL/6J wildtype male and female mice at postnatal day 3 using a previously published and well-established protocol [19]. In summary, mMG were collected using a gentle shake-off method. The cells were left in culture for an additional 24 hours before use. All experiments were conducted in DMEM supplemented with 10% fetal calf serum, 1% Pen/Strep, 1% sodium pyruvate, and 4.5 g/l d-glucose (“complete medium”).

### Primary human microglia (hMG)

Primary human microglia (hMG) were isolated from surgery tissue of patients with medication-resistant epilepsy after prior written consent. The ethics committee of the Charité Universitätsmedizin Berlin (EA2/111/14) approved all procedures. The age and sex of the patients included in this study are given in Suppl. Table 1. The microglia were isolated according to previously published protocols with slight modifications [20]. In summary, blood vessels and meninges were carefully removed from resected brain tissue using a dissection microscope (Zeiss Stemi DV4, Oberkochen, Germany). The tissue was then weighed, diced, and dissociated with the Neural Tissue Dissociation Kit (P) (#130–092-628, Miltenyi Biotec, Bergisch Gladbach, Germany). The resulting microglia were cultured as adherent cells in poly-L-lysine coated T75 flasks with microglia culture medium, which included DMEM/F12, 10% fetal calf serum, and 1X penicillin–streptomycin solution. After 1 week of incubation at 37°C and 5% CO2, cells were transferred to multiwell plates for further experiments. To detach the cells, the culture medium was removed, and the flasks were washed with phosphate-buffered saline (PBS, without Mg++ and Ca++). Cells were then incubated with 3 ml Trypsin/EDTA solution (0.25%/0.02%) at 37°C and 5% CO2 for 5 minutes. To ensure complete detachment, cells were gently scraped using a rubber cell scraper. After counting with a hemocytometer, cells were seeded in microglia culture medium on a 24-well plate.

### Induced pluripotent stem-cell derived microglia-like cells (hiPSC-MGL)

Human induced pluripotent stem cells (iPSC; cell line BIHi250-A, see https://hpscreg.eu/cell-line/BIHi250) were provided by the Berlin Institute of Health (BIH) Core Unit pluripotent Stem Cells and Organoids (CUSCO). hiPSCs were then differentiated into hiPSC-derived microglia-like cells (MGL) using a previously published protocol [17, 21]. In brief, hiPSCs were initially differentiated into hematopoietic progenitor cells over 11 days using a hematopoietic differentiation medium (HDM). From day 0 to day 4 cells were maintained in STEMdiff^TM^ hematopoietic Differentiation Medium A. From day 4 until day 10 STEMdiff^TM^ Heamotpoietic Differnetiation Medium B was applied. On day 10, cells were harvested and FACsorted based on CD43, CD45 and CD 34 expression. CD43+/CD45+ cells were then transferred to microglia differentiation medium (MDM) composed of phenol-free DMEM/F12 (1:1), ITS-G (2% v/v), B27 (2% v/v), N2 (0.5% v/v), monothioglycerol (400 μM), Glutamax (1X), NEAA (1X), and additional insulin (5 μg/ml) for another 28 days. MDM was supplemented as follows: from days 10 to 35 with M-CSF (25 ng/ml), IL-34 (100 ng/ml), and TGFb-1 (50 ng/ml); and from days 35 to 38 with M-CSF (25 ng/ml), IL-34 (100 ng/ml), TGFb-1 (50 ng/ml), CD200 (100 ng/ml), and CX3CL1 (100 ng/ml). After 38 days of differentiation, hiPSC-MGL were harvested and plated for downstream analysis.

For electrophysiological recordings cells were plated on poly-L-lysin for mMG and hMG or vitronectin for hiPSC-MGL coated glass cover slips in a 24-well plate at a density of 10.000 cells/well. For immunological stainings cells were plated in 12-well chamber slides at a density of 5.000 cells/well. For qPCR, cells were plated in 96-well plates at a density of 20.000 cells/well (mMG), 30.000 cells/well (hMG), or 40.000 cells/well (hiPSC-MGL). Cells were then incubated overnight at 37°C and 5% CO2 to allow for attachment and adaptation before stimulation with either 1 µg/ml LPS, 200 ng/ml IFN-γ, or both for 24 hours.

### Patch-clamp recordings

Cover slides with cells were placed in a custom-made patch clamp setup. Bright field images were registered using an upright monocular phototube (Leica, Bensheim, Germany) and a CCD camera (8-bit, Sanyo, Osaka, Japan) at 60X magnification.

Cells were submerged in carbogenated artificial cerebrospinal fluid (aCSF) (129 mM NaCl, 1.25 mM NaH_2_PO_4_, 1.6 mM CaCl_2_, 3 mM KCl, 1.8 mM MgSO_4_, 21 mM NaHCO_3_ and 10 mM Glucose; adjusted to 300-310 mOsm; all by Carl Roth GmbH, Karlsruhe, Germany) at room temperature. Intracellular solution consisted of 120 mM KCl, 5 mM NaCl, 2 mM MgCl_2_, 1 mM CaCl_2_, 10 mM HEPES, 11 mM EGTA (adjusted to pH 7,3 with KOH, 290-300 mOsm). Channel antagonists ML133 (20 µM) and 4-AP (1 mM) were washed in during patch clamp recordings as indicated.

Borosilicate pipettes (Science Products, Hofheim, Germany; 1.5 mm outer diameter) were pulled using a Narishige PC-10 vertical puller (London, UK; electrode resistance of 4-6 MOhm). Pipettes were placed on an Axon Instruments headstage CV-7B (sample rate 100 kHz, low-pass filter 10 kHz (8-pole Bessel Filter)), transmitting the signal to Axon CAN MultiClamp 700B (Axon Instruments, Union City, CA, USA). Signals were digitized through the Axon Digidata 1550 (Axon Instruments). Data was processed with the Multiclamp and pClamp 10 software. Cells were clamped at either −20 mV or −70 mV holding potential (as indicated in the legends). Voltage steps of 50 ms duration were applied from −170 mV to +70 mV with increasing steps of 10 mV.

### Isolation of mRNA and qPCR

RNA was extracted from cultured cells using the NucleoSpin® RNA XS kit (Macherey-Nagel, Düren, Germany). For reverse transcription, M-MLV reverse transcriptase and random hexamers were employed to convert RNA into cDNA. Polymerase chain reaction (PCR) amplification was carried out using gene-specific primers and Light Cycler® 480 SYBR Green I Master (Roche Diagnostics, Indianapolis, IN, USA). The PCR conditions were as follows: preincubation at 95°C for 10 minutes, followed by 45 cycles of 95°C for 10 seconds, primer-specific annealing temperature for 10 seconds, and 72°C for 15 seconds. Crossing points of the amplified products were determined using the Second Derivative Maximum Method (Light Cycler Version LCS480 1.5.0.39, Roche). mRNA expression levels were quantified relative to tripeptidyl peptidase 2 (*TPP2*). The specificity of the PCR products was confirmed by melting curve analysis. PCR products were also analyzed on a 1.5% agarose gel to verify the presence of a single amplicon of the expected size. Negative controls, including reactions lacking either template DNA or reverse transcriptase, showed no bands on the gel.

### Immunocytochemistry

mMG, hiPSC-MGL, and hMG were fixed for 20 minutes at room temperature using 4% paraformaldehyde (PFA). The cells were then washed three times with 1× tris-buffered saline (TBS). Non-specific binding was blocked with TBS blocking buffer (TBS+), consisting of 1× TBS, 3% normal donkey serum (NDS), and 0.1% Triton X-100 (stock solution: 10%) for 30 minutes. Primary antibodies in TBS+ were incubated overnight at 4°C. Secondary antibody was diluted in 1× TBS and species-specific donkey anti-species serum. After washing with 1x TBS, secondary antibodies were applied at a dilution of 1:400 and incubated for two hours. The cells were then washed again, and Hoechst (1.4 µg/mL), diluted in 1× TBS, was used for nuclear staining.

### Confocal microscopy

Microscopy imaging was performed using the confocal laser scanning microscope LSM 700 (Zeiss) equipped with a 40×, 1.3 NA oil objective. Representative images were obtained from random regions of the cultures as *z*-stacks with dimensions of 1,024 × 1,024 pixels in *xy*, 4 *z*-planes, and a pixel size of 0.312 × 0.312 µm.

### Image processing and morphological analysis

The morphological analysis was performed using a customized macro script in ImageJ (version 1.54f)[22]. The script applies filters to enhance the signal-to-background ratio and then creates a maximum intensity projection. The automatic thresholding algorithm “Li” was applied, followed by a series of binary operations and size filtering to fill holes and filter out small background objects. Touching cells were automatically separated by applying a watershed based on a seed image created by thresholding a distance map image using functions from the 3D ImageJ Suite plugin [23]. Objects crossing the image borders were automatically excluded. Errors in automatic segmentation were corrected manually by separating unseparated objects, excluding overlapping or merged cells with obscure borders, and manually labeling faint cellular parts that were not detected automatically. Morphological features were then extracted using the MorphoLibJ plugin and the skeleton analysis functions of ImageJ [24]. Statistical analysis was performed on selected representative features, e.g., perimeter-to-area ratio, circularity, maximum calliper diameter.

### Statistical analysis

Electrophysiological data were analyzed using custom written Matlab (Version 2021b, MathWorks, Natick, MA, USA) scripts. I/V curves were derived from the last 10 ms of each voltage pulse. Capacitance was derived from I/V curves. Inward current density was calculated at −120 mV to −100 mV and related to the cell capacitance.

Statistical analysis was performed using GraphPad Prism Version 10.2.1 (GraphPad, San Diego, CA, USA). Only for the morphological analysis, statistical testing war performed in Python using Pingouin and scikit-posthocs libraries [25, 26]. Values are represented as box plot with median, interquartile range whiskers from minimum to maximum. For the morphology analysis the whiskers were chosen to indicate 1.5x IQR, as the groups were sufficiently large. Each dot represents one measurement or cell, as indicated in the legends. The different treatment groups were compared using the Kruskal-Wallis test followed by Dunn’s test for multiple comparisons, unless indicated otherwise in the legends. Differences were estimated as significant for p<0.05 and indicated with an asterisk.

Graphical representation and figures were assorted using CorelDRAW Graphics Suits X7 (Corel Corporation, Ottawa, Canada).

## Results

### hiPSC-MGL display homeostatic microglia markers and respond to stimulation

mMG, hMG and hiPSC-MGL were cultured and characterized by immunocytochemistry and cytokine transcription to evaluate their microglial identity and validate their suitability for subsequent investigations. mMG were isolated from the brains of postnatal day 3 C57BL/6 mice. hMG were isolated from resected brain tissue obtained from patients undergoing epilepsy surgery. Only tissue from non-pathological regions was used to reflect healthy human microglia. hiPSCs were differentiated into hematopoietic progenitor cells, followed by further differentiation into MGL. All microglia populations, including mMG, hMG, and hiPSC-MGL, were analyzed for the expression of classical microglial markers, the transmembrane protein 119 (TMEM119), C-X3-C motif chemokine receptor 1 (CX3CR1), and protein tyrosine phosphatase receptor type c (CD45). Positive immunostaining for these markers was observed across all microglia groups, confirming their microglial phenotype, as previously shown (Fig. 1B, Fig. 3A)[17, 27].

**Figure 1:**
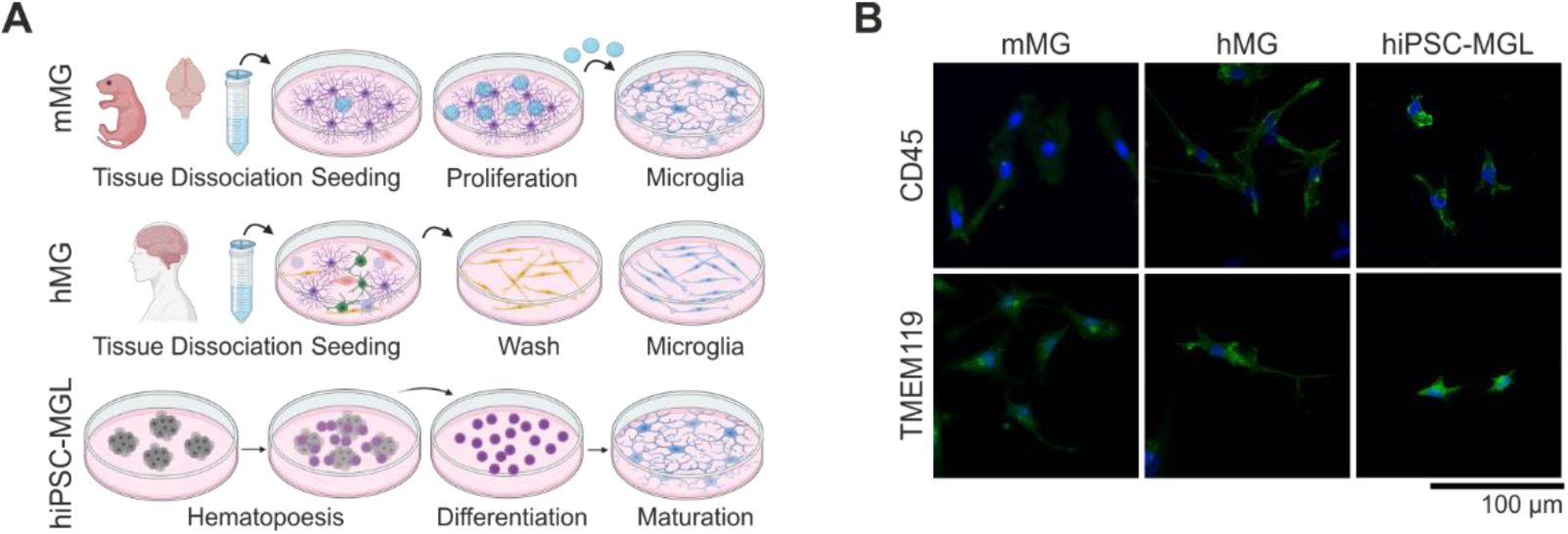
mMG, hMG and hiPSC-MGL express microglial markers. **(A)** Schematic of different sources of primary mouse microglia (mMG), primary human microglia (hMG), and human induced pluripotent stem cell-derived microglia-like cells (hiPSC-MGL). Created in BioRender. Göttert, R (2025) **(B)** Maximum intensity projections of confocal laser scanning microscopy images for mMG, hMG, and hiPSC-MGL, labeled for microglial maturations markers CD45 or TMEM119 (green) and Hoechst (blue). Scale bar 100 µm.

To assess microglial reactivity, cells were stimulated for 24 hours with lipopolysaccharide (LPS), interferon-gamma (IFN-γ), or a combination of both (LPS+IFN-γ), a protocol known to induce a robust proinflammatory phenotype in rodent microglia [28]. LPS and IFN-γ signaling pathways converge on the transcription factors nuclear factor kappa B (NFκB) and signal transducer and activator of transcription 1 (STAT1), resulting in an amplified inflammatory response [29]. As stimulation of microglia leads to the transcription of cytokines, we used quantitative real-time PCR (qPCR) to quantify messenger RNA (mRNA) expression levels of the proinflammatory cytokines tumor necrosis factor α (*TNFα*) and interleukin 6 (*IL6*), the transcription factor interferon regulatory factor 1 (*IRF1*), and the chemokine C-X-C motif chemokine ligand 11 (*CXCL11*) [30]. All microglial cell lines exhibited comparable stimulus-dependent proinflammatory responses to LPS and/or IFN-γ, with the strongest responses observed under combined stimulation conditions across all cell types (Fig. S1).

The results demonstrate that mMG and hMG retain their distinct microglial identity *in vitro*, as shown by the consistent expression of key microglial markers. Importantly, hiPSC-MGL exhibited robust expression of microglial markers, indicating successful differentiation. Furthermore, all cells expressed proinflammatory cytokines, indicating their responsiveness to the presented stimuli.

### Patch-clamp recordings show only few alterations of cell capacity in hMG and hiPSC-MGL after stimulation

The functional state of microglia was assessed using single-cell patch-clamp recordings following stimulation with proinflammatory agents. Recordings were performed in voltage clamp and membrane properties were derived from the measured IV-curves (Fig. 2A).

**Figure 2:**
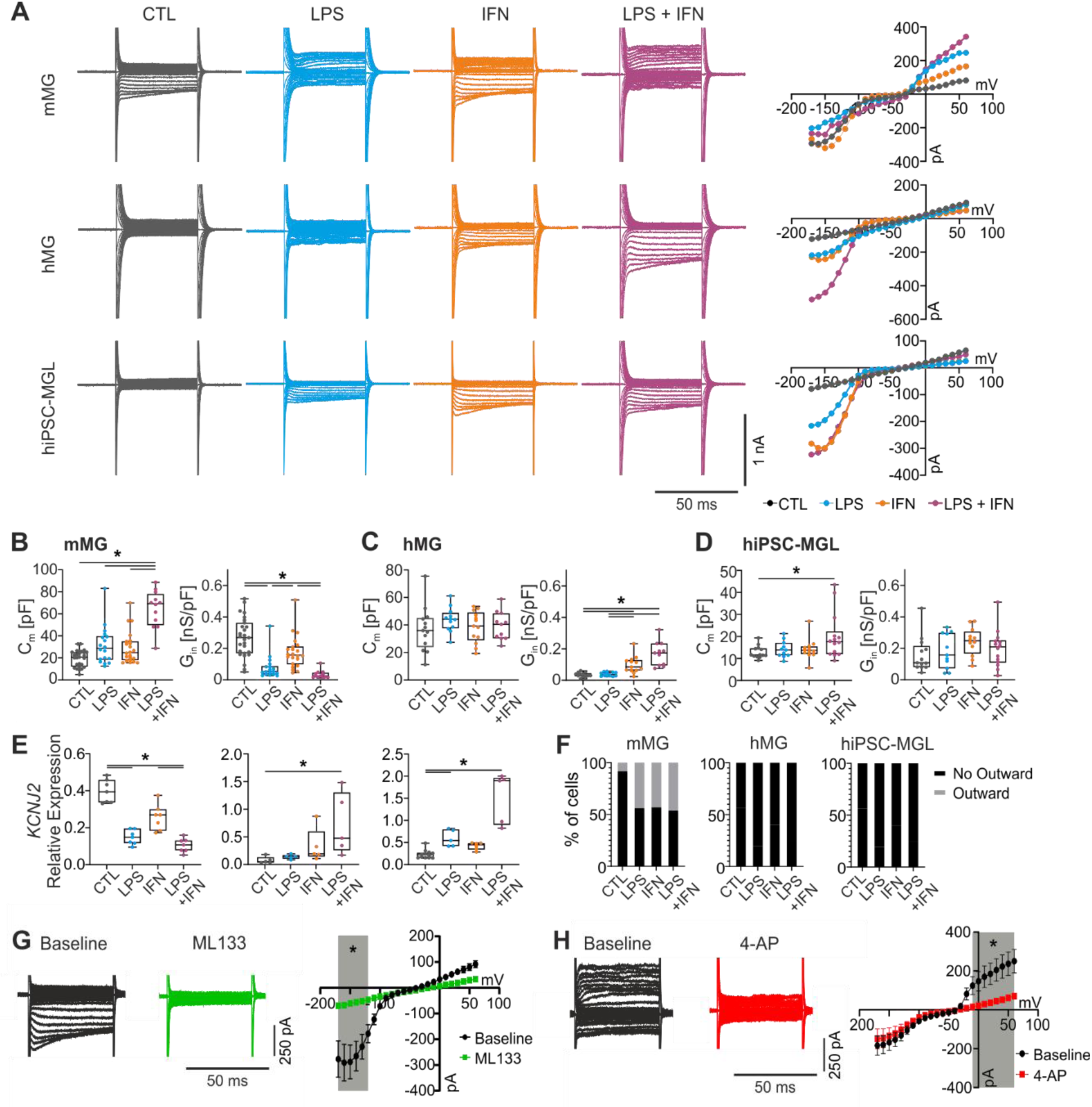
K_ir_2.1 is differently regulated in hMG and hiPSC-MGL than mMG, with no detection of K_v_ current in hMG or hiPSC-MGL. **(A)** Single-cell patch-clamp recordings of microglial cells were performed following the indicated stimulation. Representative traces of recordings at a holding potential of −70 mV of mMG, hMG, and hiPSC-MGL are shown with their corresponding IV-curves. **(B-D)** Membrane capacitance (C_m_) and inward current density (G_in_) was derived from IV-curve of whole cell patch clamp recordings at a holding potential of −20 mV. Please note the different scaling for capacitance. Statistical analysis with Kruskal-Wallis test followed by Dunn’s test for multiple comparisons, *\*P* < 0.05. Data represented as box plots with median, each dot represents one cell. For n=recordings: mMG (CTL=14, LPS=10, IFN=12, LPS+IFN=12), iPSC-MGL (CTL=13, LPS=12, IFN=12, LPS+IFN=10), hMG (CTL=11, LPS=11, IFN=12, LPS+IFN=11) **(E)** mRNA expression of *KCNJ2* was quantified by qPCR and is represented relative to householding gene *TPP2*. For n=samples: mMG (CTL=7, LPS=7, IFN=7, LPS+IFN=7), iPSC-MGL (CTL=12, LPS=5, IFN=6, LPS+IFN=5), hMG (CTL=5, LPS=5, IFN=5, LPS+IFN=5) **(F)** Percentage of cells exhibiting voltage-dependent, delayed outward current at a holding potential of −70 mV and activation at stimulation of −20 mV. N as for D-F. **(G)** Patch clamp recordings were performed before (black) and after (green) wash-in of specific K_ir_2.1 antagonist ML133 (holding potential −20 mV) in primary mouse microglia. **(H)** Patch clamp recordings were performed before (black) and after (green) wash-in of K_v_ antagonist 4-AP (holding potential −70 mV) in primary mouse microglia. IV-curves are represented as mean and standard error of the mean. Statistical analysis with two-way ANOVA and Sidak’s test for multiple comparisons, grey area indicates *\*P* < 0.05. Statistical analysis with Wilcoxon test, *\*P* < 0.05. n=8.

In mMG, stimulation with LPS+IFN-γ resulted in a significant increase in cell capacitance compared to all other groups (Fig. 2B). hMG were the most homogeneous group in terms of cell capacitance (Fig. 2C). hiPSC-MGL exhibited an increase in capacitance following LPS+IFN-γ (Fig. 2D). Overall, hiPSC-MGL displayed lower capacitance than mMG and hMG (note different scale). In summary, mMG demonstrated distinct electrophysiological changes regarding cell capacitance, while the changes of hMG and hiPSC-MGL were less pronounced.

### hMG but not mMG increase K_ir_2.1 currents upon activation with LPS and IFN-γ

Next, we performed a detailed analysis to characterize specific microglial currents, starting with an inward current with time-dependent inactivation. For mMG, this inward current displayed the highest current density under control conditions, with a significant downregulation observed following stimulation with LPS alone and in combination with IFN-γ. Stimulation with IFN-γ did not significantly alter the current density (Fig. 2B). Microglia isolated from adult mice also displayed a reduction in inward current density following stimulation with LPS+IFN-γ (Fig. S2D). Inversely, hMG showed a significant enhancement of the inward current following IFN-γ stimulation, with an even more pronounced increase observed after combined LPS+IFN-γ treatment (Fig. 2C). In hiPSC-MGL, stimulation with LPS and/or IFN-γ did not result in a significant increase in inward current conductivity; however, there was a general trend towards stronger currents following proinflammatory stimulation (Fig. 2D).

The inward current’s characteristics of time-dependent inactivation at low voltages suggested the involvement of the inward rectifying potassium channel K_ir_2.1 (Fig. S2A) [11]. To investigate this, we performed qPCR of mRNA levels of the potassium inwardly rectifying channel subfamily J member 2 (*KCNJ2*), the gene encoding the K_ir_2.1 channel. In mMG stimulation with LPS and LPS+IFN-γ led to a significant downregulation of *Kcnj2* expression. In hMG a significant increase in *KCNJ2* expression was observed following combined LPS+IFN-γ stimulation. In hiPSC-MGL LPS and LPS+IFN-γ stimulation resulted in a significant upregulation of *KCNJ2* transcripts (Fig. 2E). These transcriptional alterations mirrored the observed changes in current density, implicating K_ir_2.1 as the most likely ion channel involved. Furthermore, RNA sequencing data from human brain samples published in the Human Protein Atlas and by Olah et al confirm the expression of *KCNJ2* in human microglia (Fig. S2F, G). To prove the role of K_ir_2.1, the specific antagonist ML133 was applied during patch-clamp recordings of mMG exhibiting the characteristic inward current. Indeed, application of ML133 completely abolished the current, supporting the hypothesis that K_ir_2.1 mediates this response (Fig. 2G).

In summary, the K_ir_2.1 channel is downregulated in mMG following stimulation with LPS and LPS+IFN-γ, correlating with a decreased inward current. Conversely, hMG and hiPSC-MGL demonstrate increased expression and functional activity of K_ir_2.1 in response to inflammatory stimuli.

### hMG and hiPSC-MGL display no K_v_ current upon stimulation

The presence of an outward current that was deactivated at a holding potential of −20 mV but could be activated at a holding potential of −70 mV and stimulation above −20 mV, exhibiting delayed activation, was evaluated next (Fig. S2B). This current was detectable in a subset of mMG under control conditions, with a notable increase following stimulation with LPS and/or IFN-γ (Fig. 2F). This current was also present in a small subset of microglia isolated from adult mice (Fig. S2E). The described current characteristics are characteristic of K_v_ channels. We confirmed this, by applying the K_v_ channel antagonist 4-AP to mMG exhibiting the delayed-outward current, which led to the current’s abolition (Fig. 2H). Remarkably, the K_v_ current was completely absent in hMG and hiPSC-MGL, both under control conditions and following the various stimulations (Fig. 2F). In line with this, data published in the Human Protein Atlas and the dataset from Olah et al. also failed to show any relevant expression of *KCNA3*, the gene encoding the prominent K_v_ channel 1.3, in human microglia under healthy or pathological conditions (Fig. S2F, G).

These findings highlight electrophysiological variability between different species. The absence of K_v_ currents in hMG and hiPSC-MGL is of particular interest, as it has been discussed as a potential drug target to reduce inflammation in mice.

### hMG and hiPSC-MGL do not display amoeboid morphology after stimulation

Microglia are known to alter their morphology as part of their response after exposure to pathological stimuli. Furthermore, ion channel activity and the resting membrane potential are important factors for morphological changes. In order to evaluate cell morphology, we visualized the cells using a CX3CR1-antibody (Fig. 3A) and performed a supervised automated morphological analysis (details see Methods Image Processing and morphological analysis).

**Figure 3:**
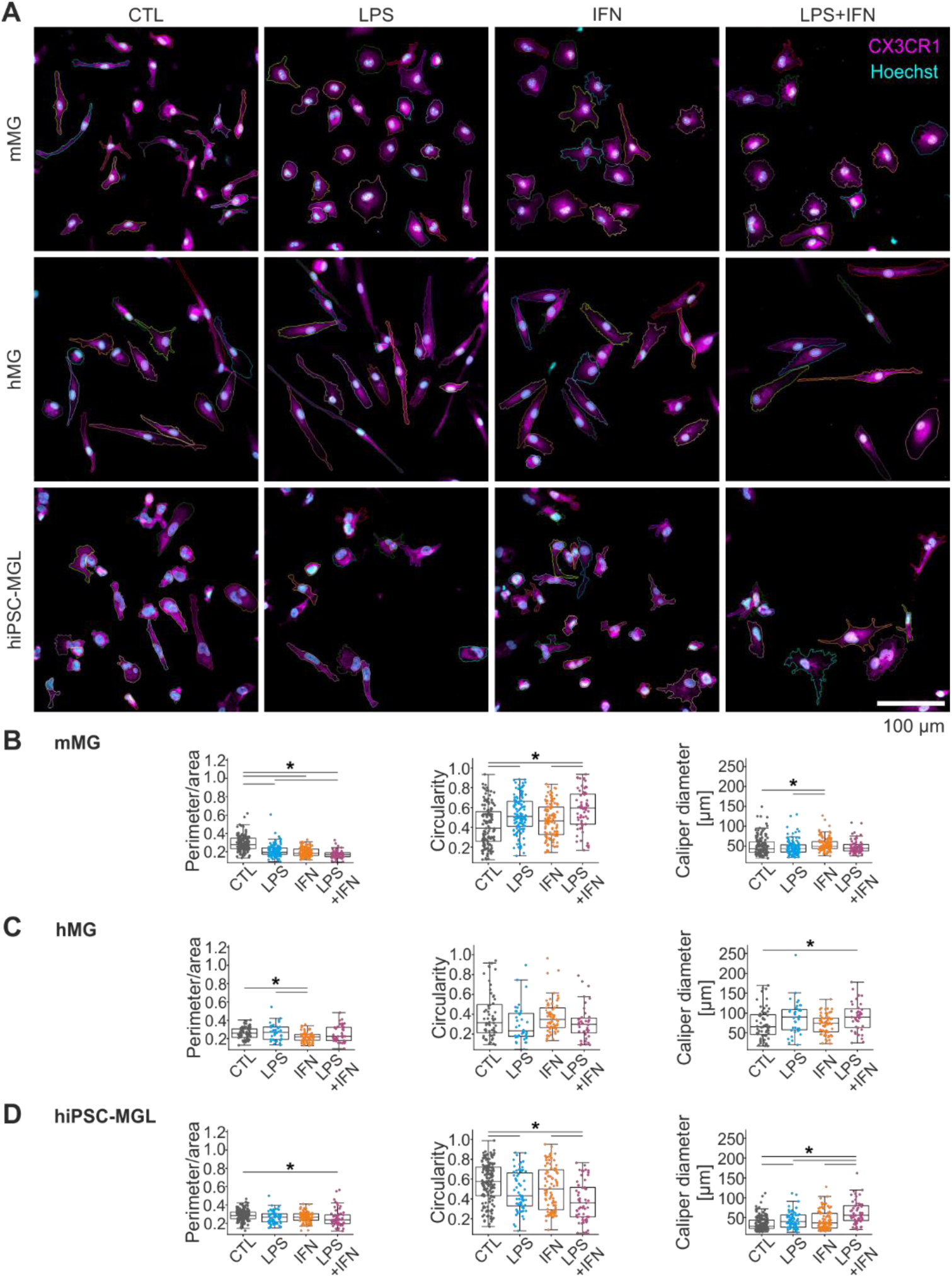
hMG show little morphological alterations, while hiPSC-MGL increase ramification after stimulation. **(A)** Maximum intensity projections of confocal laser scanning microscopy images for cultured cells from the studied cell lines under four treatment conditions, labeled with CX3CR1-AF488 (magenta) and Hoechst (cyan). The colored borders of cells and nuclei represent the segmentation results of the morphological analysis. Cells were excluded if they crossed the image borders, overlapped with other cells, or merged with other cells with obscure borders. Scale bar 100 µm. **(B-D)** Statistical analysis of several morphological features quantifying the morphological changes of cultured cells under the treatment conditions. Statistical analysis with Kruskal-Wallis test followed by Dunn’s test for multiple comparisons, *\*P* < 0.05. Data represented as box plot with median. Each dot represents one cell. For n=cells: mMG (CTL=135, LPS=123, IFN=107, LPS+IFN=63), iPSC-MGL (CTL=154, LPS=63, IFN=88, LPS+IFN=57), hMG (CTL=60, LPS=38, IFN=61, LPS+IFN=39).

In mMG, LPS increased cell circularity, with even greater effects observed under LPS+IFN-γ stimulation. IFN-γ alone increased maximum caliper diameter. All stimuli led to a reduction in the perimeter-to-area ratio, indicating a loss of ramification and shift towards an amoeboid morphology in mMG (Fig. 3B). This reflects the electrophysiological increase in cell capacitance (Fig. 2B). hMG exhibited a more elongated morphology at baseline and only few morphological changes after stimulation. IFN-γ induced a slight decrease in perimeter/area, and LPS+IFN-γ led to a modest increase in caliper diameter, with no major alterations in circularity (Fig. 3C). In hiPSC-MGL, LPS and LPS+IFN-γ decreased circularity and increased caliper diameter, while LPS+IFN-γ also decreased perimeter/area (Fig. 3D). As numbers of end-point voxels were also increased, this indicates a more ramified morphology after stimulation (Fig. S3D).

Overall, mMG displayed a more ameboid morphology following stimulation with LPS+IFN-γ, whereas hMG did not show strong morphological alterations. In contrast, hiPSC-MGL reacted with increased ramification to the stimulation.

## Discussion

Microglia express a range of ion channels which are correlated to different functional states and are being evaluated as potential drug targets. However, microglial ion channel activity has mostly been studied in rodent microglia with little information being available from human microglia. Geirsdottir et al. showed that microglia express core genes across species, however notable differences were found in several gene modules between rodent and human microglia, raising the question regarding the translation of mouse data to the human situation [31]. Differences between human and murine microglial sensosome have been described particularly in the context of pathology, constituting a particular translational challenge [32].

Therefore, in this study, we investigated the functional response of human microglia, using hMG and hiPSC-MGL, in established stimulation protocols and compared them with mMG. Our work yielded the following major results: first, we did not see K_v_ currents in hMG; second, K_ir_2.1 is regulated differently in hMG than in mMG; and third, hiPSC-MGL exhibit similar electrophysiological properties to hMG.

We showed the presence of an outward-rectifying potassium current in mMG which was increased after stimulation, in mMG isolated from both postnatal and adult mice. Notably, this current was absent in hMG and hiPSC-MGL under all stimulation conditions. The outward current’s dynamics and sensitivity to 4-AP confirmed its classification within the voltage-gated K_v_ family. This is consistent with previous evidence identifying these currents as K_v_1.3 and/or K_v_1.5 mediated [33, 34]. K_v_1.3 is of particular interest, as its activity has been linked to induction of a microglial proinflammatory state [33, 35]. However, K_v_1.3 studies employing patch-clamp techniques have mainly been performed in animal models or immortalized rodent cell lines such as BV2 [36, 37]. Studies have identified K_v_1.3 as a potential microglial drug target in various neurological diseases, such as Parkinson’s disease, stroke or Alzheimer’s disease [14, 38, 39]. While K_v_1.3 expression has been demonstrated in human T-cells and pulmonary macrophages, electrophysiological evidence for K_v_1.3 in human microglia remains lacking [40–42]. Reports of detection of K_v_1.3 in human microglia rely primarily on indirect detection methods (e.g., Western blot, immunocytochemistry, or PCR), leaving a gap in functional data [15, 38, 39]. Furthermore, evidence is inconsistent, as single-cell sequencing and the human protein atlas show little to no microglial *KCNA3* transcription [43]. The data of Olah et al. also indicate no clear upregulation of *KCNA3* in an aged human population or in Alzheimer’s disease patients [44, 45]. Our data suggest that the K_v_1.3 channel may not be active in human microglia. Furthermore, the intact cytokine transcription suggests a K_v_-independent activation of a proinflammatory state in hMG. Similarly, Nguyen et al. previously described a lack of K_v_ current in patch-clamp recordings of cultured human microglia [46]. Interestingly, the comprehensive proteome database of human and mouse microglia of Lloyed et al. shows a lack of K_v_1.3 in both hMG and human embryonic stem-cell derived (hESC)-MGL, whereas it could be detected in xenografted-hESC-MGL to mice, suggesting a context-dependent expression in human microglia [47]. The discrepancy in K_v_1.3 activity in human and mouse microglia could partly explain the translational failure of K_v_1.3-targeting microglial therapies and highlights the need to carefully consider species- and context-dependent channel expression.

Consistent with prior reports in rodent microglia, we found that mMG exhibited a prominent inward-rectifying potassium current (K_ir_2.1) at baseline [9, 11]. Interestingly, this current is typically absent in rodent microglia *in vivo* unless exposed to pathogenic stimuli, suggesting that *in vitro* isolation procedures may inadvertently induce a pre-activated state in mMG [9]. Contrary to *in vivo*, K_ir_2.1 current density decreased in mMG from both postnatal and adult mice following LPS+IFN-γ stimulation. This goes in line with the depolarized reversal potential, as K_ir_2.1 contributes to stabilizing the resting membrane potential. In hMG, however, stimulation led to an increase in K_ir_2.1 currents, resembling the activation profile observed in mouse microglia *in vivo* [4]. This high expression of K_ir_2.1 can be correlated to the retained elongated microglial morphology of hMG, even after stimulation. This correlation was even stronger in hiPSC-MGL. The membrane potential determines the driving force for calcium entry, linking hyperpolarization to rapid stimulus–response coupling and a more ramified microglial morphology [34, 48]. In the dataset published by Olah et al the expression of *KCNJ2* is mainly upregulated in clusters associated with an anti-inflammatory profile (Fig. S2G) [44]. Meanwhile, the antagonism of K_ir_2.1 has also been shown to reduce proliferation and migration in rodent microglia [11, 49]. These findings suggest different regulatory mechanisms of K_ir_2.1 in human and mouse microglial responses to proinflammatory signals *in vitro*. Furthermore, the lack of amoeboid morphology following stimulation of human microglia correlates with K_ir_2.1 activity. Further studies are necessary to better understand the specific mechanistic function of K_ir_2.1 in morphology and inflammatory response.

We demonstrated that hiPSC-MGL exhibited robust expression of microglial markers, indicating successful differentiation, comparable to primary microglia. These findings validate the use of hiPSC-MGL as a reliable model for further mechanistic studies, as previously shown [6, 17]. We also showed that hiPSC-MGL were responsive to the stimulation with LPS and/or IFN-γ, as it led to an increase in transcription of proinflammatory cytokines, which was largely consistent across cell types and in line with existing literature [18, 50, 51]. We could show that single-cell patch-clamp recordings were not just feasible but also hold a lot of potential for hiPSC-MGL. Furthermore, they more closely mirrored hMG reactivity to inflammatory stimuli than mMG, exhibiting no K_v_ currents while showing a tendency toward increased K_ir_2.1 activity upon stimulation. Interestingly, they showed an increased ramification after stimulation. Overall, these changes go in line with the described alterations in hMG, and in opposition to mMG. Functional studies on hMG remain rare due to limited tissue availability and technical challenges in *in situ* labeling, further underscoring the value of hiPSC-MGL [52]. The use of hMG and hiPSC-MGL allowed for a significant reduction in animals needed for this study. Although animal-derived supplements were still used in this study, the models offer the possibility of working completely animal-free, such as by using recombinant proteins from HEK cells [53, 54]. This reduces the reliance on animal experiments and the ethical concerns associated with animal use.

In summary, our study underscores critical differences between mouse and human microglia in terms of electrophysiological properties, ion channel expression as a potential drug target, and morphology. Although mMG remain a widely used model system, our data highlight important species-specific differences that might explain translational inconsistencies. The use of hiPSC-MGL is a promising bridging technology and reduces animal numbers in scientific experiments. Incorporating human-derived microglia into preclinical pipelines may therefore help bridge the translational gap between experimental findings and clinical outcomes.

### Limitations of the study

A possible limitation of this study is the varied cell age, as microglial phenotype and receptor expression are known to vary across developmental stages and with aging [55]. mMG were isolated at postnatal day 3 as this constitutes a well-established reference system, widely used in numerous studies and forming the basis for further investigations, particularly in pathological contexts. Culturing adult mouse microglia entails several known technical limitations, including low yield and purity, and reduced viability [56, 57]. However, we could show similar ion current patterns in postnatal and adult mouse microglia. Furthermore, it is known that microglia from adult rodents show strong K_v_ and K_ir_2.1 currents following pathological stimulation *in vivo* [4, 9]. The hiPSC-MGLs originated from reprogrammed fibroblasts of adult donors, and hMG were isolated from adult surgical specimens. This heterogeneity in donor age may contribute to differences in microglial responses; however, the most pronounced effects regarding potassium current presence were observed between murine and human cells, regardless of age.

Additionally, differences in culture conditions could have influenced cellular behavior. The medium used for hiPSC-MGLs contained macrophage colony-stimulating factor (M-CSF), which is known to enhance microglial proliferation and phagocytic activity [58]. IL-34, also contained in the MGL culture medium, has been associated with the promotion of a more mature and neuroprotective microglial phenotype [59]. In contrast, mMG and hMG were not exposed to M-CSF or IL-34. Both mMG and hMG incubation media included fetal calf serum, however, which has previously been reported to induce microglial phagocytic activity [60, 61]. While these supplements are standard in microglial differentiation protocols, they may also contribute to variability in functional response to stimuli.

## Resource Availability

### Lead contact

Requests for further information and resources should be directed to and will be fulfilled by the corresponding author, Karen Gertz.

### Material availability

This study did not generate new unique reagents.

### Data and code availability

All data reported in this paper will be shared by the lead contact upon request. This paper does not report original code.

Any additional information requires to reanalyze the data reported in this paper is available from the lead contact upon request.

## Acknowledgment

The authors would like to acknowledge the support of the Berlin Institute of Health Core Unit pluripotent Stem Cells and Organoids (CUSCO) for providing the hiPSC line. This study was supported by the Deutsche Forschungsgemeinschaft (Priority Program 2395/GE, 2576/6-1 to K.G.; Germanýs Excellence Strategy—EXC-2049—390688087 to M.E.; Collaborative Research Center ReTune TRR 295-424778381 to M.E.; Clinical Research Group KFO 5023 BeCAUSE-Y, project 2 EN343/16-1 to M.E.), the Bundesministerium für Bildung und Forschung (CSB to M.E., and K.G.), the German Center for Neurodegenerative Diseases (DZNE to M.E.), the German Center for Cardiovascular Research (DZHK to M.E. and K.G.), the German Center for Mental Health (DZPG to M.E.). We would like to thank Jan-Oliver Hollnagel in particular for his support in creating the MATLAB script for analyzing the electrophysiological data. Special thanks go to Melanie Kroh, Stefanie Balz and Bettina Herrmann for their excellent technical assistance.

## Author contribution

S.S., R.G., and K.G. conceived this study. S.S. performed the electrophysiological experiments. J.F. and R.G. performed cell culturing, immunostaining and microscopy. S.S. analyzed the data and M.K. performed the morphological analysis. A.P. and J.H. collected human brain tissue. J.O., T.S., T.K., M.S. performed neurosurgical procedures. S.S., R.G. and K.G. wrote the manuscript with input from all authors. All authors discussed the results and commented on the final manuscript.

## Resource Table

**Table 1:** Agents.

| REAGENT or RESOURCE | SOURCE | IDENTIFIER |
| --- | --- | --- |
| **Antibodies** |  |  |
| Anti-CD45 (1:100) | Abd Serotec,<br>Neuried, Germany | Cat # MCA1031GA<br>RRID: AB_324339 |
| Anti-CD45 (human, 1:100) | EXBIO, Praha,<br>Czech Republic | Cat # 11-222-C025<br>RRID: AB_10732878 |
| Anti-CX3CR1 (human, 1:50) | Bio-Rad,<br>Feldkirchen,<br>Germany | Cat # AHP1589<br>RRID: AB_2087421 |
| Anti-TMEM119 (1:100) | ATLAS AK,<br>Bromma, Sweden | Cat # HPA051870<br>RRID: AB_2681645 |
| Anti-mouse (Alexa 488, 1:400) | Invitrogen,<br>Waltham, MA, USA | Cat # A21202<br>RRID: AB_141607 |
| Anti-rat (Alexa 488, 1:400) | Invitrogen,<br>Waltham, MA, USA | Cat # A21208<br>RRID: AB_2535794 |
| Anti-rabbit (Alexa 488, 1:400) | Invitrogen,<br>Waltham, MA, USA | Cat # A21206<br>RRID: AB_2535792 |
| PE anti-human CD45 Antibody (1:20) | BioLegend, San<br>Diego, CA, USA | Cat # 304007<br>RRID: AB_314395 |
| APC anti-human CD43 Antibody (1:20) | BioLegend, San<br>Diego, CA, USA | Cat # 343205<br>RRID: AB_2194073 |
| FITC anti-human CD34 Antibody | BioLegend, San<br>Diego, CA, USA | Cat # 343603<br>RRID: AB_1732030 |
| **Bacterial and virus strains** |  |  |
| None |  |  |
| **Biological samples** |  |  |
| None |  |  |
| **Chemicals, peptides, and recombinant proteins** |  |  |
| 4-AP | Sigma-Aldrich, St.<br>Louis, MO, USA | Cat # 275875-5G |
| ML133 | Sigma-Aldrich, St.<br>Louis, MO, USA | Cat # SML0190-5MG |
| LPS from E. coli 055:B5 | Sigma-Aldrich, St.<br>Louis, MO, USA | Cat # L4524-10MG |
| Human IFN-g | Peprtech,<br>Cranbury, NJ, USA | Cat # 300-02 |
| Murine IFN-g | Peprtech,<br>Cranbury, NJ, USA | Cat # 315-05 |
| DMEM | Thermo Fisher,<br>Waltham, MA, USA | Cat # 31966-021 |
| FCS | PAN Biotech<br>Aidenbach,<br>Germany | Cat # P30-3031 |
| Pen/Strep | PAN Biotech,<br>Aidenbach,<br>Germany | Cat # P06-07100 |
| Sodium pyruvate | PAN Biotech<br>Aidenbach,<br>Germany | Cat # P04-43100 |
| D-Glucose | Sigma-Aldrich, St.<br>Louis, MO, USA | Cat # G8769-100ML |
| STEMdiff | Stemcell<br>Technologies,<br>Vancouver, Canada | Cat # 05311 |
| DMEM/F12 | Thermo Fisher,<br>Waltham, MA, USA | Cat # 11039-021 |
| ITS-G | Thermo Fisher,<br>Waltham, MA, USA | Cat # 41400-045 |
| B27 | Thermo Fisher,<br>Waltham, MA, USA | Cat # 17504-044 |
| N-2 | Thermo Fisher,<br>Waltham, MA, USA | Cat # 17502-048 |
| Monothioglycerol | Sigma-Aldrich, St.<br>Louis, MO, USA | Cat # M1753-100ML |
| Glutamax | Thermo Fisher,<br>Waltham, MA, USA | Cat # 35050-061 |
| NEAA | Thermo Fisher,<br>Waltham, MA, USA | Cat # 11140-050 |
| Insulin | PAN Biotech,<br>Aidenbach,<br>Germany | Cat # P07-04300 |
| M-CSF | Peprtech,<br>Cranbury, NJ, USA | Cat # 300-25 |
| IL-34 | Peprtech,<br>Cranbury, NJ, USA | Cat # 200-34 |
| TGFb1 | Peprtech,<br>Cranbury, NJ, USA | Cat # 100-21 |
| CD200 | Novoprotein,<br>Shanghai, China | Cat # C311 |
| CX3CL1 | Peprtech,<br>Cranbury, NJ, USA | Cat # 300-31-50UG |
| PBS (no Mg, No Ca) | Thermo Fisher,<br>Waltham, MA, USA | Cat # 14200-067 |
| Trypsin/EDTA | PAN Biotech,<br>Aidenbach,<br>Germany | Cat # P10-024100 |
| PLL | Sigma-Aldrich, St.<br>Louis, MO, USA | Cat # P4707 |
| Vitronectin | Thermo Fisher,<br>Waltham, MA, USA | Cat # A14700 |
| Matrigel | Corning Incorporated,<br>Corning, NY, USA | Cat # 354277 |
| M-MLV reverse transcriptase | Promega, Madison, WI, Germany | Cat # M1701 |
| Random hexamers | Promega, Madison, WI, Germany | Cat # C1181 |
| Agarose gel | Carl Roth GmbH, Karlsruhe, Germany | Cat # 2267.4 |
| Paraformaldehyd | Carl Roth GmbH, Karlsruhe, Germany | Cat # 0335.2 |
| Donkey Serum | Dominique Dutscher, Bernolsheim, France | Cat # S2170-100 |
| Triton X-100 | Sigma-Aldrich, St. Louis, MO, USA | Cat # X100-100ML |
| Hoechst | Abcam, Cambridge, UK | Cat # 33342 |
| **Critical commercial assays** |  |  |
| Nucleo Spin RNA kit | Macherey-Nagel, Düren, Germany | Cat # 740902.50 |
| **Deposited data** |  |  |
| None |  |  |
| **Experimental models: Cell lines** |  |  |
| BIHi250-A | BIH, Berlin, Germany | https://hpscereg.eu/cell-line/BIHi250-A |
| **Experimental models: Organisms/strains** |  |  |
| C57BL/6J wildtype mice |  |  |
| **Oligonucleotides** |  |  |
| None |  |  |
| **Recombinant DNA** |  |  |
| None |  |  |
| **Software and algorithms** |  |  |
| pClamp 10 | Molecular Devices, San Jose, CA, USA |  |
| Matlab (Version 2021b,) | MathWorks, Natick, MA, USA |  |
| GraphPad Prism Version 10.2.1 | GraphPad, San Diego, CA, USA |  |
| CorelDRAW Graphics Suits X7 | Corel Corporation, Ottawa, Canada |  |
| **Other** |  |  |
| Cell scraper | SPL Life Sciences, Pocheon-si, South Korea | Cat # 90021 |
| T75 flask | Corning Incorporated,<br>Corning, NY, USA | Cat # 353136 |
| Glass cover slips, 5 mm | Assistant,<br>Glaswarenfabrik<br>Karl Hecht,<br>Sondheim vor der<br>Rhön, Germany | Cat # 41001 |
| 12 well chamber slides | Ibidi, Gräfeling,<br>Germany | Cat # 81201 |

**Table 2:**
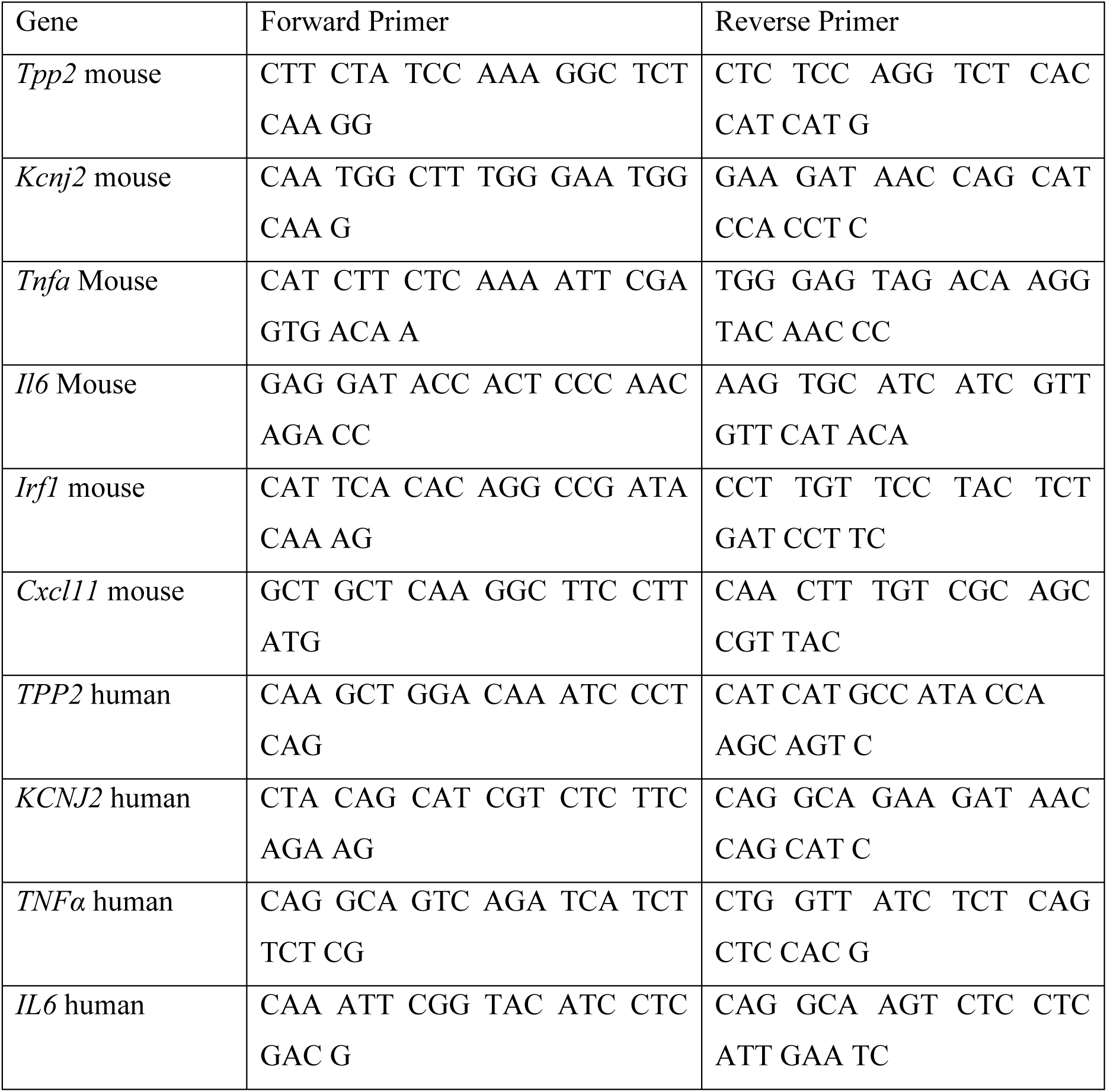
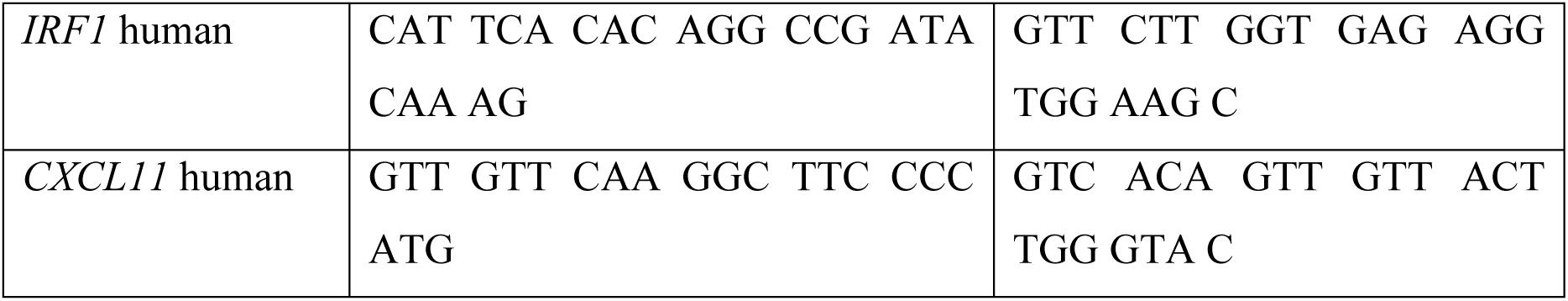
Primer sequences.

