## Supplementary Figures for "Mind the translational gap: human microglia differ from mouse microglia in their regulation of K_v_ and K_ir_2.1 channels"

### Supplemental information

#### Material and Methods

Primary mouse microglia from adult mice (14–16 weeks old) were isolated from whole brains without the cerebellum, as previously described, with slight modifications [S1,S2]. In brief, blood vessels and meninges were carefully removed from the resected brain tissue. The tissue was then diced and dissociated using the Neural Tissue Dissociation Kit (P) (#130–092-628, Miltenyi Biotec, Bergisch Gladbach, Germany). Microglia obtained from one mouse brain were cultured as adherent cells in non-coated T25 flasks containing microglia culture medium composed of DMEM/F12 supplemented with with 10% fetal calf serum and 1X penicillin–streptomycin solution. One day after the isolation procedure, the medium was supplemented with macrophage colony-stimulating factor (M-CSF, 100 ng/mL, PeproTech #315-02) and granulocyte-macrophage colony-stimulating factor (GM-CSF, 100 ng/mL, PeproTech #315-03). Microglia were cultured for 7 days, and the medium was changed every 3 days. After 1 week of incubation at 37°C and 5% $CO_2$, the cells were transferred to PLL-coated glass coverslips for patch-clamp experiments. To detach the cells, the culture medium was removed, and the flasks were washed with phosphate-buffered saline (PBS, without $Mg^{2+}$ and $Ca^{2+}$). The cells were then incubated with 0.5 mM EDTA solution (diluted from 0.5 M solution, Invitrogen, #15575020) at 37°C and 5% $CO_2$ for 10 minutes. To ensure complete detachment, cells were gently scraped using a rubber cell scraper and seeded in microglia culture medium without M-CSF and GM-CSF.

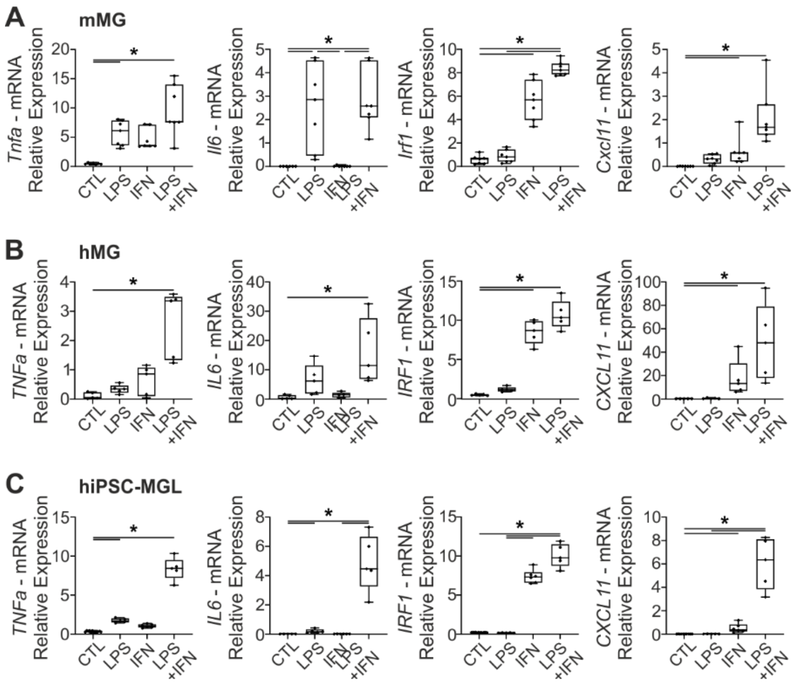


**Fig. S1: mMG, hMG, and hiPSC-MGL upregulate cytokines upon proinflammatory stimulation**

**B.** mRNA of *TNFa, IL6, IRF1* and *CXCL11* were quantified by qPCR and are presented relative to the mRNA expression of householding gene *TPP2*. Cells were collected after stimulation with the indicated stimuli.

Statistical analysis with the Kruskal-Wallis test followed by Dunn's test for multiple comparisons, **P* < 0.05. Data represented box plots and median. Each dot represents one sample.

For n=samples: mMG (CTL n=7, LPS n=7, IFN n=7, LPS+IFN n=7), iPSC-MGL (CTL n=12, LPS n=5, IFN n=6, LPS+IFN n=5), hMG (CTL n=5, LPS n=5, IFN n=5, LPS+IFN n=5)

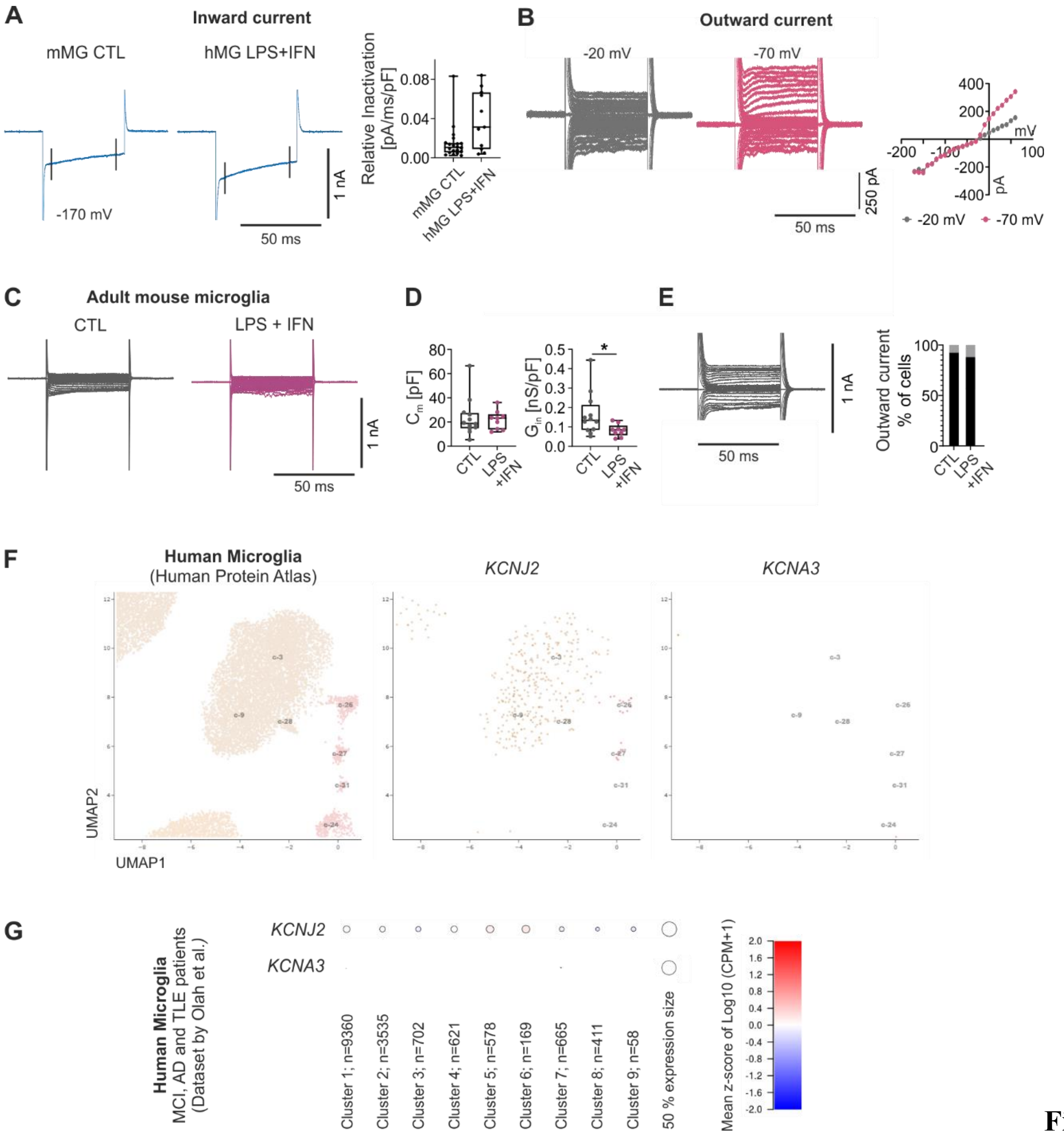


**Fig. S2: $K_v$ and $K_{ir}2.1$ characteristics, currents in adult mouse microglia and expression in human tissue**

**(A)** Time-dependent inactivation of inward currents at low voltages. Exemplary traces at -170 mV and a holding potential of -20 mV are shown for mMG under control condition and hMG after stimulation with LPS+IFN-γ. The inactivation is calculated from the current at the indicated time points (black bars) and the decrease over time. The values are normalized to the capacity of each cell. For n=recodings: mMG CTL n=20, hMG LPS+IFN n=11 (B) Exemplary recording of mMG after stimulation with LPS+IFN-γ displaying voltage-dependent delayed outward rectifying current and corresponding IV-curves. At a holding potential of -20 mV (grey) the channel is inactive, whereas at a holding potential of -70 mV (pink) it displays the characteristic delayed inward current. **(C)** Single-cell patch-clamp recordings of primary mouse microglia isolated from adult mice were performed following stimulation with LPS and IFN-γ. Representative traces of recordings at a holding potential of -20 mV are shown. **(D)** The basic membrane characteristics of reversal potential ($V_{rev}$), membrane resistance ($R_m$), capacitance ($C_m$) and specific inward conductance ($G_{in}$) were derived from the IV-curves. Statistical analysis with the Mann-Whitney test, $^*P < 0.05$. Data represented as box plots with median, each dot represents one cell. **(E)** Percentage of cells exhibiting voltage-dependent, delayed outward current at a holding potential of -70 mV and activation at stimulation of -20 mV with exemplary trace. For n=recordings: CTL n=12, LPS+IFN n=9. **(F)** UMAP from single cell analysis in the human protein atlas (www.proteinatlas.org). Microglia are represented in Cluster c-3, c-9, and c-28. *KCNJ2* had the following expression in the three microglia clusters: c-3 189 reads/4332 cells, Expression 11.9 nCPM; c-9 155 reads/2534 cells, Expression 27.9 nCPM, c-28 9 reads/133 cells, Expression 12.7 nCPM. *KCNA3* had no reads in any of the three microglia clusters. **(G)** RNA expression derived from the dataset of Olah et al. [1]. Microglia were isolated from postmortem tissue from patients suffering from Alzheimer's disease (AD), mild cognitive impairment (MCI) or from fresh surgery tissue of patients suffering from temporal lobe epilepsy (TLE). Based on their expression profiles, the microglia were then grouped into 9 different clusters. The data show induced expression of *KCNJ2* in clusters 5 and 6. Cluster 5 is marked by increased interleukin 10 signaling and both clusters 5 and 6 show increased expression of CD83, overall relating clusters 5 and 6 to an anti-inflammatory response. There is no relevant expression of *KCNA3* in any of the clusters.

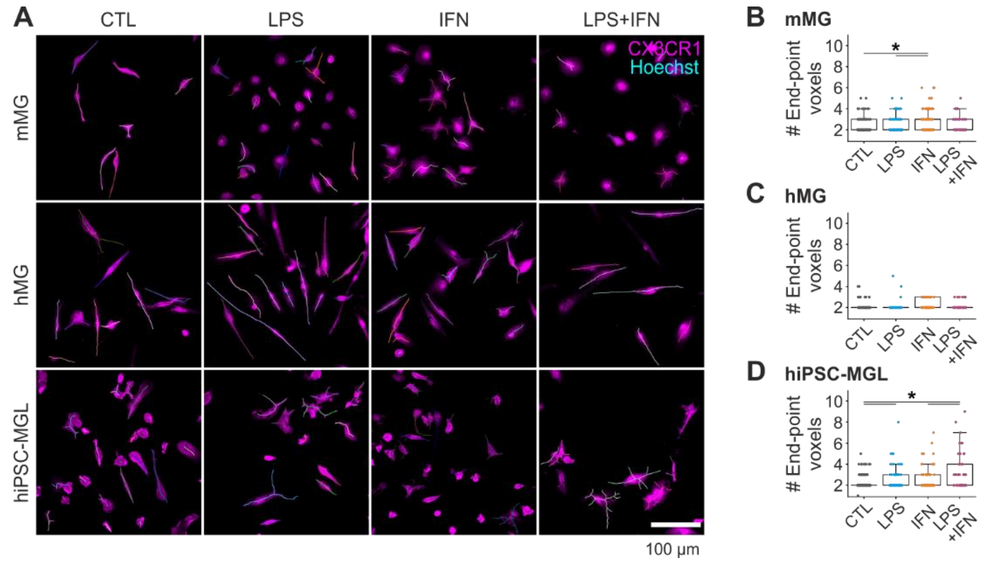


**Fig. S3: Analysis of bifurcation of microglial cell cultures**

**(A)** Maximum intensity projections of confocal laser scanning microscopy images for cultured cells from the studied cell lines under four treatment conditions, labeled with CX3CR1-AF488 (magenta) and Hoechst (cyan). The colored lines represent the skeleton results of the morphological analysis. Scale bar: 100 µm. **(B-E)** Statistical analysis of number of End-point Voxels, quantifying the ramification of cultured cells under the treatment conditions. Statistical analysis with the Kruskal-Wallis test followed by Dunn's test for multiple comparisons, $^{*}P < 0.05$. Data represented as box plot with median. Each dot represents one cell.

For n=cells: mMG (CTL n=135, LPS n=123, IFN n=107, LPS+IFN n=63), iPSC-MGL (CTL n=154, LPS n=63, IFN n=88, LPS+IFN n=57), hMG (CTL n=60, LPS n=38, IFN n=61, LPS+IFN n=39).

**Suppl. Table 1**: **Characteristics of human donors**

| ID# | Sex | Age | Pathology |
|---|---|---|---|
| Pat #1 | male | 55 | TLE |
| Pat #2 | male | 59 | TLE |
| Pat #5 | male | 40 | TLE |
| Pat #6 | male | 39 | TLE |
| Pat #7 | male | 29 | TLE |
| Pat #9 | female | 37 | TLE |
| Pat #14 | male | 58 | TLE |
| hiPSC | Female | - | |
